# Chromosome-level assembly of the common vetch reference genome (*Vicia sativa*)

**DOI:** 10.1101/2021.10.11.464017

**Authors:** Hangwei Xi, Vy Nguyen, Christopher Ward, Zhipeng Liu, Iain R. Searle

## Abstract

**Background:** *Vicia sativa* L. (Common Vetch, n = 6) is an annual, herbaceous, climbing legume that is distributed in tropical, sub-tropical and temperate climates. Originating in the Fertile Crescent of the Middle East, *V. sativa* is now widespread and grows in the Mediterranean basin, West, Central and Eastern Asia, North and South America. *V. sativa* is of economic importance as a forage legume in countries such as Australia, China, and the USA and contributes valuable nitrogen to agricultural rotation cropping systems. To accelerate precision genome breeding and genomics-based selection of this legume, we here present a chromosome-level reference genome sequence for *V. sativa*.

**Results:** We applied a combination of long-read Oxford Nanopore sequencing, short-read Illumina sequencing, and high-throughput chromosome conformation data (CHiCAGO and Hi-C) analysis to construct a chromosome-level genome of *V. sativa*. The chromosome-level assembly of six pseudo-chromosomes has a total genome length of 1.9 gigabases (Gb) with a median contig length of 684 kb. Benchmarking Universal Single-Copy Orthologs (BUSCO) of the assembly demonstrated a very high completeness of 98 % of the dicotyledonous orthologs. RNA-seq analysis and gene modelling enabled the annotation of 58,415 protein-coding genes.

**Conclusions:** The high-quality chromosome-level genome assembly of *V. sativa* will provide novel insights into vetch genome evolution and be a valuable resource for genomic breeding, genetic diversity and for understanding adaption to diverse arid environments.

## Background

*Vicia sativa* L. (common vetch) (Fig. 1) is an annual legume belonging to the *Fabaceae* family, and *Vicia* genus [1]. The *Vicia* genus contains about 180 - 210 species including the economically important crop broad bean [2]. To date, no draft or chromosome-level genome assembly within the *Vicia* genus has been reported. Interestingly, *V. sativa* has at least three different reported haploid chromosome numbers, n = 5, 6 or 7 [3] while n= 6 is the best characterized karyotype.

**Figure 1:**
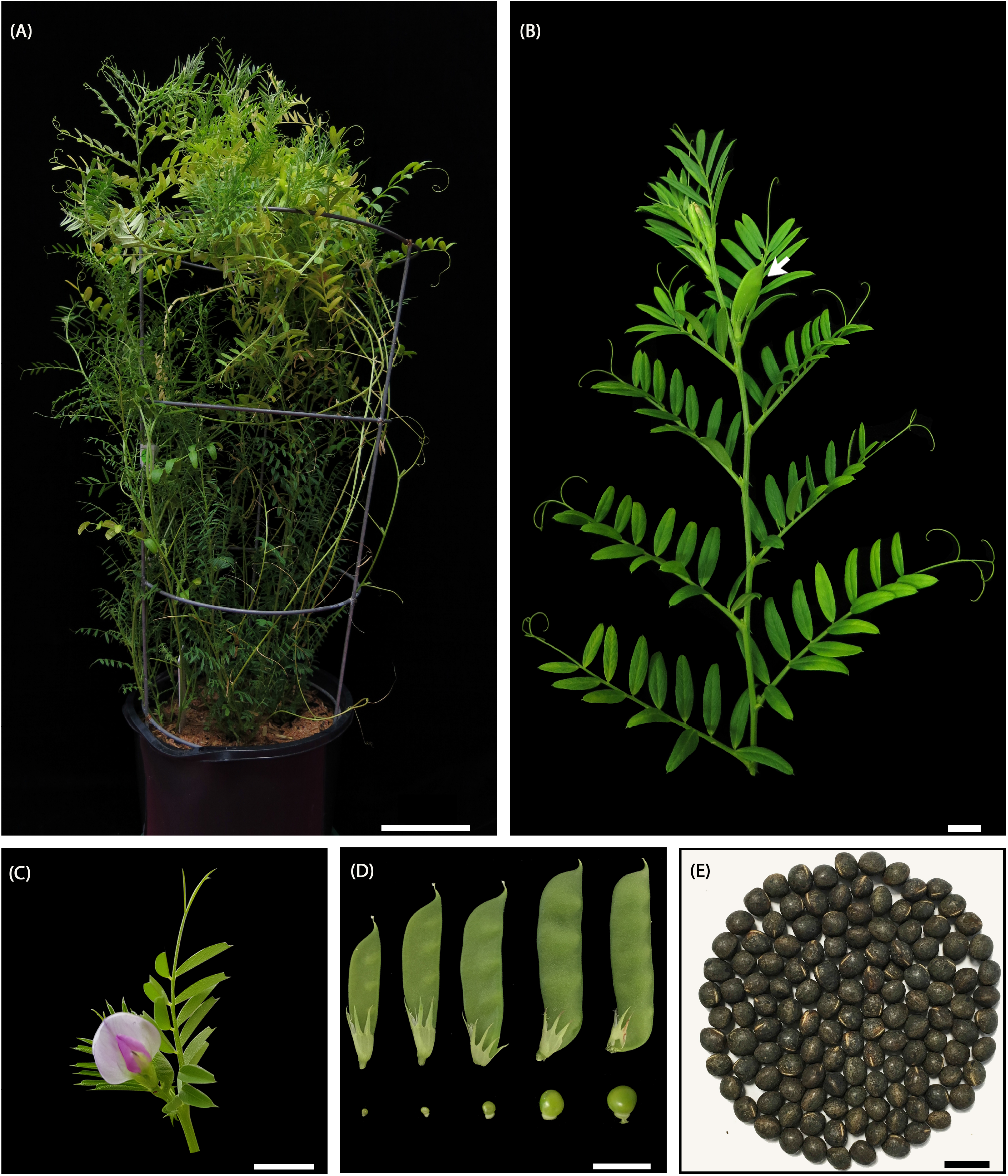
*V. sativa* (cultivar Studenica). (A) Glasshouse grown ten-week-old *V. sativa* at flowering. (B) Detached stem showing compound leaves at each node. At the end of each compound leaf is a tendril. A single pod forms at the base of each leaf node after flowering (arrow). (C) A shoot apex with a flower and surrounding young leaves. (D) Young to mature seed pods (left to right), with a representative seed shown at the bottom of each pod. Each pod contains three to five seeds. (E)Dry seeds of cultivar Studenica. Scale bars (A) = 10 cm, (B, C, E) = 1 cm, (D) = 2 cm.

*V. sativa* is thought to have originated in the Fertile Crescent of the Middle East and is now widespread on every continent as both a crop and a weed [4]. *V. sativa* is a multi-purpose legume, and the plants are often grown for forage and the seeds can be used safely as a feed for ruminant animals. *V. sativa* seed contains up to 30 % crude protein and is rich in essential amino acids and unsaturated fatty acids [5]. However, only a small amount of the seed can be safely fed to monogastric animals, examples include chickens and pigs, due to the presence of neurotoxic proteinaceous amino acids β-cyano-L-alanine and γ-glutamyl-β-cyano-alanine [6].

*V. sativa* is often used in crop-rotation systems to increase nitrogen input to the soil. A study of *V. sativa* - wheat rotation in four-year-period showed that the cultivation of *V. sativa* during autumn increased soil water storage and subsequently increased biological yield and grain yield of wheat, in which both yields were doubled in the third year compared to the second year of the rotation [7]. Furthermore, the symbiosis between soil *rhizobia* bacteria with *V. sativa* roots allows the plant to fix atmospheric nitrogen and later provide nitrogen for the following crop, hence reducing the use of expensive nitrogen fertilizer [8]. *V. sativa* exhibits an excellent drought tolerance and is suitable to be cultivated in arid areas. In one drought resistance study, *V. sativa* could withstand a month of drought stress, with the leaf weight not decreasing significantly compared to the non-drought control [9]. *V. sativa* offers multiple-usage and is a valuable crop in a sustainable agricultural system [10].

With the important value of *V. sativa*, vetch breeders have primarily selected for traits conferring high yield, pod shattering, flowering time, disease resistance against *Ascochyta fabae, Uromyces viciae-fabae* (rust) and *Sclerotinia sclerotium* [11]. Recently published transcriptome data has facilitated the uncovering of agricultural important traits at the gene expression level, such as pod-shattering resistance [12] and drought tolerance genes [13] in *V. sativa*. However, a lack of high-quality genome reference is currently impeding the genetic mapping of important genes and hindering further applications such as genome editing when compared to other crops.

In this study, we assembled a high-quality chromosome level reference genome for *V. sativa* which is the first chromosome-level reference genome in the *Vicia* genus. We performed genome annotation using RNA-seq data from five tissues to ensure capturing most of the expressed genes. We also included a phylogenetic analysis of *V. sativa* and legume relatives. We envisage that our *V. sativa* genome will be an important resource for the evolutionary studies of this species. The well annotated chromosome-level genome will also provide important information to facilitate genetic mapping, gene discovery and functional gene studies.

## Data description

### Sampling and sequencing

To prepare *V. sativa* whole genome sequencing (WGS) long-read and shortread data, seeds of cultivar Studenica (*V. sativa* subsp. *sativa*) was obtained from The South Australian Research and Development Institute (SARDI, South Australia, Australia). Seeds were sterilised and germinated *in vitro* on ½ Murashige & Skoog basal medium with 1 % sucrose for three days at 25 °C, in the dark. Bulk three-millimetre-long primary root tips were then harvested and snap-frozen in liquid nitrogen for subsequent DNA extraction. DNA was extracted using phenol:chloroform method [14], with an additional high-salt low-ethanol wash to improve DNA purity [15]. High quality DNA was confirmed by electrophoresis on 1 % agarose gel. The DNA was sent to the Australian Genome Research Facility (AGRF, Melbourne, Australia), and Novogene Co., Ltd (HongKong, China) for library preparation and sequencing on a PromethION (PromethION, <u>RRID:SCR_017987</u>) and Novo-Seq 6000 (Illumina NovaSeq 6000 Sequencing System, <u>RRID:SCR_016387</u>), respectively. We obtained 72 Gb of Nanopore long-read data, and 205 Gb paired-end short-read data (150 bp read length).

To produce *V. sativa* CHiCAGO sequencing data [16] and Hi-C sequencing data [17], two grams of young leaf tissue was snap-frozen in liquid nitrogen and sent to Dovetail Genomics (USA) for library preparation and sequencing. CHiCAGO and Hi-C libraries were sequenced on an Illumina HiSeq X (Illumina HiSeq X Ten, <u>RRID:SCR_016385</u>) to produce 162 Gb of CHiCAGO and 148 Gb of Hi-C sequencing data, respectively.

To prepare *V. sativa* RNA sequencing (RNA-seq) data, RNA was purified from the first two fully expanded leaves, shoot apexes with young leaves up to 1 cm long from 4-week old plants, roots from 5-day old seedlings and 4-week old leaf-derived callus tissues using the Spectrum™ Plant Total RNA Kit (Sigma Aldrich) with additional DNase I treatment to remove DNA contamination (On-Column DNase I Digestion, Sigma Aldrich), and ribosome removal treatment to enrich for non-ribosomal RNA fraction (Ribo-Zero rRNA Removal Kit for Plant Leaf or Plant Seed/Root, Illumina) [18]. Directional RNA libraries were prepared for each tissue using the NEBNext Ultra™ Directional RNA Library Prep Kit for Illumina (New England Biolabs) following the manufacturer’s protocol. Libraries were sent to Novogene Co., Ltd (HongKong, China) for sequencing on Novo-Seq 6000 (Illumina) to obtain 150 bp paired-end read data. In total, we obtained 74.6 Gb of RNA-seq data. A summary of the long and short-read sequencing data is provided in Table 1.

**Table 1:** Overview of sequencing data generated in this study.

| Libraries | Insert size (bp) | Raw data (Gb) | Clean data (Gb) | Mean read length (bp) | Coverage (×)* |
| --- | --- | --- | --- | --- | --- |
| WGS Illumina short-reads | 300 | 205.13 | 200.28 | 150 | 106.28 |
| Nanopore reads | N/A | 72.12 | N/A | 9,094 | 37.38 |
| Chicago | 350 | 162.00 | N/A | 150 | 83.94 |
| Hi-C | 350 | 147.60 | N/A | 150 | 76.47 |
| Illumina RNA-seq reads | 300 | 74.60 | 66.49 | 150 | 38.65 |

\*Coverage was calculated based on the assembled genome size of 1.93 Gb.

### Genome size estimation and genome assembly

We first performed a genome size estimation for *V. sativa*. To do this, resequenced short-reads were trimmed using Trim Galore v0.4.2 (Trim Galore, <u>RRID:SCR_011847</u>) with default parameters and 25-mers were counted using Jellyfish v2.2.6 (Jellyfish, <u>RRID:SCR_005491</u>) [19]. The 25-mer count distribution data was used as an input to Genomescope (GenomeScope, <u>RRID:SCR_017014</u>) [20] for genome size estimation with the read length set to 150 and max k-mer coverage set to 1 million. Genomescope estimated a genome wide heterozygosity level of 0.09 % (Supplementary Fig. S1) and a genome size of 1.61 Gb, approximately 160 Mb smaller than the genome size estimated by the flow cytometry (1.77 Gb) [21].

Next, we conducted contig assembly from the Nanopore long-reads using Canu v1.7 (Canu, <u>RRID:SCR_015880</u>) [22] under default parameters with the expected genome size set at 1.77 Gb. Canu was used to perform read trimming and sequencing error correction for the input data before performing contig assembly. The assembled contigs were polished using clean WGS short-reads with Pilon v1.22 (Pilon, <u>RRID:SCR_014731</u>) [23] under default parameters. We repeated the polishing step and observed a further improvement of the contig quality (Supplementary Table S1). The contig quality was assessed using BUSCO v5.2.2 (BUSCO, <u>RRID:SCR_015008</u>) [24] for the completeness of the genome, and after two rounds of polishing, the complete BUSCOs increased from 69.9 % to 97.8 % (Supplementary Table S1). Using this polished assembly, a very high proportion, 99.8 %, of the WGS short-read data mapped to this final assembly (Supplementary Table S2). Overall, we obtained 9,990 assembled contigs which were 1.93 Gb, with a N50 value of 685 kb, and a mean GC content of 35 % (Table 2).

**Table 2:** Overview of *V. sativa* genome assembly.

| Feature | Value |
| --- | --- |
| Total length | 1,933,655,782 bp |
| No. of contigs | 9,990 |
| N50 length (contig) | 684,593 bp |
| N50 length (scaffold) | 290,126,875 bp |
| GC content | 35.02 % |
| Predicted protein-coding genes | 58,330 |
| Predicted noncoding genes | 10,149 |
| Content of repetitive sequences | 84.37 % |

### Chromosome-level assembly using Hi-C and linkage map data

To generate a chromosome-level assembly for *V. sativa*, Hi-C proximity [25] ligation data and CHiCAGO [26] were used to order and orient the contigs along chromosomes. The scaffolding process was carried out by Dovetail Genomics (Santa Cruz, CA, USA) using the Dovetail™ Hi-C library reads to connect and order the input set of contigs. After scaffolding with HiRise (v2.1.7) [51], the assembled genome sequence initially comprised a total of 1.8 Gb, with a scaffold and contig N50 of 51.4 and 0.6 Mb, respectively. A high fraction of the assembled sequences (93 %) were contained in four pseudo-chromosomes (Supplementary Fig. S2A), however *V. sativa* has six pairs of chromosomes [1]. We observed two of the four pseudo-chromosomes had weak interactions suggesting mis-joining of two contigs (Supplementary Fig. S2A). In parallel to the HiRise analysis, we performed a second chromosome-level assembly using 3D-DNA (3D de novo assembly, <u>RRID:SCR_017227</u>) [27].

3D-DNA scaffolding was performed by first mapping the Hi-C reads to the contig assembly using Juicer v1.6 (Juicer, <u>RRID:SCR_017226</u>) [28], which then generated 304,484,352 uniquely mapped pair-end reads and of which 51.1 % (155,477,211) of the uniquely mapped reads were identified as valid Hi-C contacts. The 3D-DNA v180114 pipeline was integrated to anchor contigs to pseudochromosomes based on valid Hi-C contacts. The output file was used to generate a Hi-C heat map for manually inspected using Juicebox Assembly Tools v1.11.08 (Juicebox, <u>RRID:SCR_021172</u>) revealing six high-quality pseudo-chromosomes (Supplementary Fig. S2A).

We compared the HiRise and 3D-DNA assembled pseudo-chromosomes by performing a whole genome alignment with Mummer v4.0.0 (MUMmer, <u>RRID:SCR_018171</u>) [29]. The alignment showed a strong synteny between the HiRise and 3D-DNA pseudo-chromosomes (Supplementary Fig. S2B). However, the two longest HiRise pseudo-chromosomes aligned to four 3D-DNA pseudo-chromosomes suggesting two HiRise pseudo-chromosomes were mis-joined (Supplementary Fig. S2B). The putative mis-joined HiRise pseudo-chromosomes also coincided with low Hi-C interactions (Supplementary Fig. S2A).

To further support that these two HiRise pseudo-chromosomes were misjoined, we compared the synteny of the HiRise and 3D-DNA pseudo-chromosomes to the high-quality *V. faba* genetic linkage map [30] as no genetic linkage map is available for *V. sativa*. When we compared the order of 1,536 sequenced *V. faba* DNA markers to their homologous regions in our HiRise and 3D-DNA pseudo-chromosomes, we observed a clear synteny between *V. faba* and *V. sativa*. However, two out of four of the HiRise pseudo-chromosomes appeared to be mis-joined, for example, the markers on HiRise pseudo-chromosomes one, mapped to two *V. faba* linkage groups (Supplementary Fig. S3). After combining the karyote, Hi-C interaction and synteny data to *V. faba*, we concluded the 3D-DNA assembly was most likely correct and subsequently used this assembly in further analysis. The final version of the genome assembly contains six pseudo-chromosomes, in which a total of 1.65 Gb contigs are anchored to these pseudo-chromosomes, representing 85.41 % of the assembled genome (Fig. 2).

**Figure 2:**
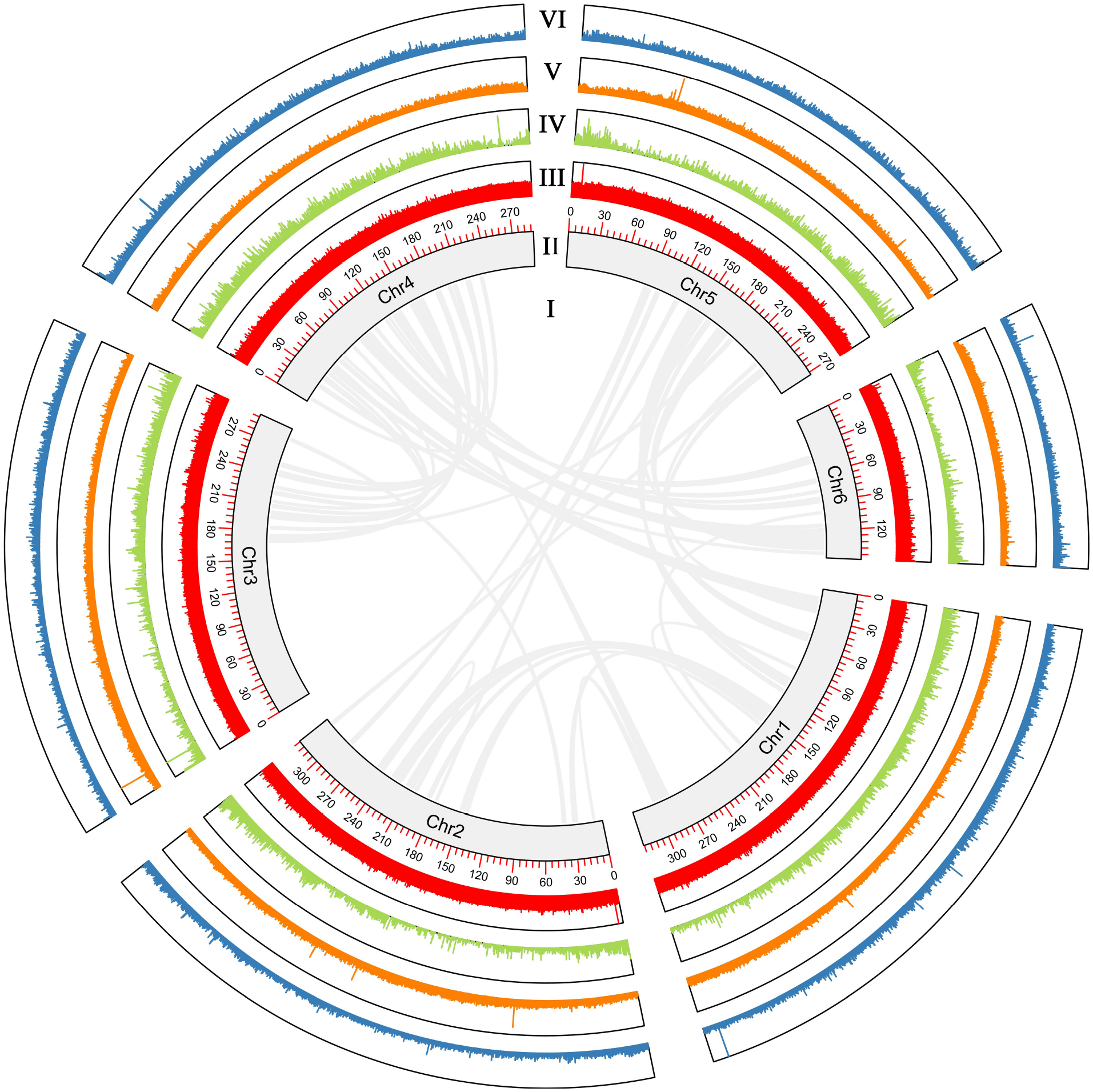
Circos plot showing the characterization of the *V. sativa* genome assembly. (I) Syntenic regions within the *V. sativa* genome based on homology searches using MCscan in Jcvi (MCScan, <u>RRID:SCR_017650</u>) [60] requiring ≥ 10 genes per block (links). (II) Pseudo-chromosome length in Mb. (III) GC content in non-overlapping 10 kb windows (histograms). (IV) Gene density in non-overlapping 10 kb windows (histograms). (V) LTR-transposable element content in non-overlapping 10 kb windows. (VI) Mutator TIR transposon content in non-overlapping 10 kb windows (histograms). Percentage of GC content, gene density, and transposable element density were calculated relative to the highest value present in the genome. Chr = pseudo-chromosome.

To assess the quality of our final version of genome assembly, genome completeness was assessed by using BUSCO v 5.2.2 referencing fabales_odb10 gene sets. Overall, BUSCO identified 97.8 % complete genes (of which 8.9 % were duplicated), 0.3 % fragmented genes, and 1.9 % missing genes out of 5,366 markers in the gene sets, suggesting the high completeness of the assembly.

### Genome annotation

To annotate the *V. sativa* genome assembly, we masked the repeat regions of the genome, then mapped the RNA-seq data to the masked genome and performed gene prediction. First, the repeat families found in the *V. sativa* genome were identified *de novo* and classified using the software package EDTA v1.9.6 [31] with the sensitive model setting. EDTA integrated multiple programs including LTR_FINDER (LTR_Finder, <u>RRID:SCR_015247</u>) [32] and RepeatModeler (RepeatModeler, <u>RRID:SCR_015027</u>), which generated a non-redundant transposable element (TE) library used to annotate the TE regions on the genome. The TE library generated from the EDTA was also used as an input to RepeatMasker v4.1.2 (RepeatMasker, <u>RRID:SCR_012954</u>) to identify and perform “hard-masking” and “soft-masking” for the repetitive region on the genome. A total of 84.37 % of the genome was masked, and 60.63% of the genome was detected as LTR elements (Supplementary Table S3).

After the genome masking, a combination of *ab initio* prediction and transcript evidence from the RNA-seq were used for gene prediction. Briefly, each RNA-seq data set was trimmed for low quality bases using Trim Galore v0.4.2, mapped to the hard-masked-genome by using STAR v2.7.9 (STAR, <u>RRID:SCR_004463</u>) [33] to generate BAM files. Then the soft-masked-genome and the BAM files generated from STAR were used to perform gene predictions using BRAKER v2.1.6 (BRAKER, <u>RRID:SCR_018964</u>) [34]. As result, a total of 58,330 predicted protein-coding-genes were identified (Supplementary Table S4). To assess the completeness of these protein-coding-genes, BUSCO v5.1.3 with fabales_odb10 gene sets were used which then identified 5,178 (96.5 %) complete, 535 (10.0 %) duplicated, 37 fragmented (0.7 %) and 151 missing (2.8 %) orthologs.

Putative functions of the predicted protein-coding-genes were characterised by comparing the predicted proteins against the Swissprot and NCBI non-redundant database using Diamond v2.0.11 (DIAMOND, <u>RRID:SCR_016071</u>) [35] with e-value cut-off of 1e-5. Protein motifs and domains were annotated by comparing the predicted proteins against the Interpro database using Interproscan v5.52-86.0 (InterProScan, <u>RRID:SCR_005829</u>) [36]. The predicted proteins were also assigned with Gene Ontology (GO) terms corresponding to the InterPro entries using Interproscan v5.52-86.0. In addition, we compared the predicted proteins against the EggNOG database v5.0 (eggnog, <u>RRID:SCR_002456)</u> [37] using eggNOG-mapper v2.1.4-2 (eggNOG-mapper, <u>RRID:SCR_021165</u>) [38] and assigned them with KEGG pathways and KEGG orthologous groups (KO). As a result, we were able to annotate a total of 51,974 (89.1 %) predicted protein coding genes with at least one function term (supplementary Table S5).

In addition, we also identified and annotated non-coding RNA in the *V. sativa* genome. tRNA was identified using tRNAscan-SE v2.07 (tRNAscan-SE, <u>RRID:SCR_010835</u>) [39], and other types of non-coding RNA were identified by using Infernal v1.1.4 (Infernal, <u>RRID:SCR_011809</u>) [40] based on the Rfam database (Rfam, <u>RRID:SCR_007891</u>) [41], overall 10,149 of noncoding genes were annotated, including 165 miRNA, 6,501 rRNA and 1,947 snRNA (Supplementary Table S6).

### Phylogenetic tree construction and divergence time estimation

We identified the orthogroups, phylogenetic position and divergence times between *V. sativa* and ten other plant species. The protein-coding sequences (Supplementary Table S7) used in our analysis were obtained from the Phytozome and GigaDB databases. First, protein sequences of *V. sativa, Medicago truncatula, Trifolium pratense, Phaseolus vulgaris, P. lunatus, Vigna unguiculata, Chamaecrista fasciculata, Faidherbia albida, Cercis canadensis, Carya illinoinensis*, and *Arabidopsis thaliana* were clustered into orthogroups using the Orthofinder v2.5.4 (OrthoFinder, <u>RRID:SCR_017118</u>) [42] under default parameters. A total of 9,122 single-copy and 49,208 multi-copy genes were identified in the *V. sativa* annotation (Fig. 3B, Supplementary Table S8), forming 19,114 orthogroups (Fig. 3A, Supplementary Table S8). Comparing orthogroups amongst *V. sativa, M. truncatula, T. pratense, P. vulgaris, F. albida*, we identified 2,654 orthogroups that are specific to *V. sativa* (Fig. 3A). Orthofinder was further used to perform phylogenetic reconstruction with the multiple sequence alignment approach (using the -msa command). The generated species tree has a support value of one on all nodes (Supplementary Figure S4), indicating the high reliability of the revealed phylogenetic relationships.

**Figure 3:**
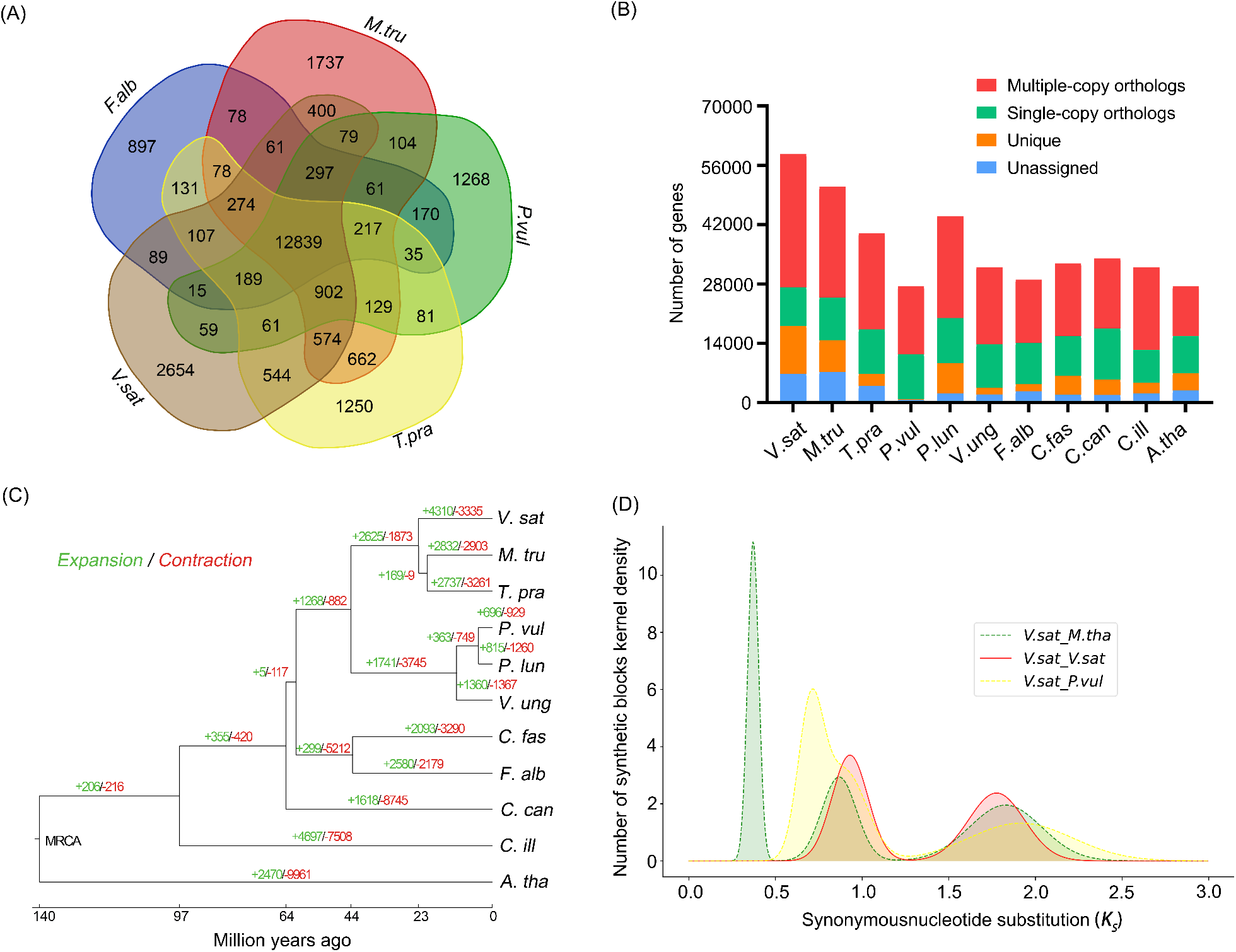
Evolution of the *V. sativa* genome. (A) A Venn diagram showing shared and unique orthologous gene families in *V. sativa* and four other legumes. (B) Predicted orthologous protein composition for *V. sativa* compared to *A. thaliana*, *C. illinoinensis* and eight legumes. (C) A phylogenetic tree shows the expansion and contraction of the gene families and the divergence time for species. (D) K_s_ plot shows the whole genome duplication event in *V. sativa, M. truncatula* and *P. vulgaris. V. sat: Vicia sativa, M. tru: Medicago truncatula, T. pra: Trifolium pratense, P. vul: Phaseolus vulgaris, P. lun: Phaseolus lunatus, V. ung: Vigna unguiculata, C. fas: Chamaecrista fasciculata, F. alb: Faidherbia albida, C. can: Cercis canadensis, C. ill: Carya illinoinensis, A. tha: Arabidopsis thaliana*.

To estimate divergence times between *V. sativa* and other important legume species (Supplementary Table S7), coding sequences of 64 randomly selected single copy orthologs were aligned using MACSE v1.2 [43]. Low-quality regions of each alignment were trimmed using Trimal v1.4.1 (trimAl, <u>RRID:SCR_017334</u>) [44], resulting in high-quality alignments totalling 95,064 bp. Individual alignments were then imported into Beast v2.6.3 (BEAST2, <u>RRID:SCR_017307</u>) [45] for phylogenetic dating. Substitution models were selected using BEAST Model Test [46] for each alignment and were allowed to coalesce using unlinked relaxed log-normal molecular clocks [47]. A calibrated Yule prior [48] was used to inform tree building and speciation with four node calibrations. First, a log normal distribution of 89.3 MYA (5 % quantile 97.9 MYA, median 106 MYA, 95 % quantile 121 MYA) [49] was used to inform the root prior (Brassicaceae, Fabaceae split). Three fossil calibrations (Supplementary Table S9) were then set using CladeAge [50]: i) Fabaceae (Supplementary Figure S5 red dot; 65.3 MYA) [51], ii) Caesalpinioideae (Supplementary Figure S5 blue dot; 58 MYA) [52, 53], and iii) Papilionoideae (Supplementary Figure S5 green dot; 55 MYA) [54]. Furthermore, a net diversification rate was set to 0.1 – 0.134 to construct a distribution around the literature value of 0.117 [55], turnover rate was set to 0.823 to 0.883 to construct a distribution around the literature value of 0.853 [55], and sampling rate of 0.000034 - 0.013 [56] were set to determine CladeAge prior distributions. Final chain length of the MCMC (288 million) was determined through continuous examination of the log file using Tracer until proper mixing was observed. This allowed us to determine a robust estimate for the MRCA of *V. sativa* and *M. truncatula* at 22.8 (95 % Highest Posterior Density: 21.6 - 24.1) MYA (Supplementary Figure S5). Gene family expansion and contraction analysis using CAFE v4.2.1 (Computational Analysis of gene Family Evolution, <u>RRID:SCR_018924</u>) [57] with a single λ revealed that there are 7,625 gene families that have undergone gene expansion or contraction since the MRCA of *V. sativa* (n = 4,310) and *M. truncatula* (n = 3,315, Fig. 3C).

To identify whole genome duplication events (WGD), WGDI v 0.5.1 [58] was used to identify gene collinearity between *V. sativa, M. truncatula* and *P. vulgaris*. The K_s_ (synonymous substitutions per synonymous site) value was calculated based on the identified collinearity gene to construct a frequency distribution map. The Ks distribution indicated that *V. sativa, M. truncatula* and *P. vulgaris* share the same ancestral WGD event. The estimated time of this WGD event (~58 MYA) [59] and the corresponding Ks value (~0.93, Fig. 3D) reveal that the average mutation rate of *V. sativa* genome is 8.02 × 10^−9^ per site per year.

## Conclusion

Understanding the genetic, epigenetic and epitranscriptomic basis of the evolutionary processes shaping drought tolerance, low nutrient requirements and adaption to broad habitats requires the comparison of multiple legume genomes, preferentially assembled at the chromosome level. In this study, we presented a complete chromosome-level genome assembly for the legume *V. sativa* (Common Vetch) and provided a detailed genome annotation. There are >19,000 species of legumes, about 200 within the *Vicia* genus, and this genome will serve as an excellent reference for the assembly of other *Vicia* genomes. The *V. sativa* genome will also facilitate comparative analyses aimed at understanding the evolutionary origin and dynamics of legume specific gene families. Our new *V. sativa* genome will greatly benefit legume researchers and plant breeders who are interested in conventional as well as engineering crop improvement.

## Availability of supporting data

Final assembly and original Nanopore assembly, as well as annotation files, predicted transcript and protein sequences, and bioinformatics supporting information, were deposited in the database GigaDB. Additionally, assembly, Illumina and Nanopore subreads, and transcriptome raw data are available in NCBI and can be accessed with BioProject PRJNA762450 and BioSample SAMN21393724. Illumina and Nanopore subreads can be obtained, with SRR16004114 and SRR16004115; and RNA-sequencing raw reads, as follows: SAMN21545804, SAMN21545805, SAMN21545806, SAMN21545807 and SAMN21545808.

## Abbreviations

CHiCAGO: Capture Hi-C Analysis of Genomic Organisation
Gb: gigabase pairs
kb: kilobase pairs
Mb: megabase pairs
MYA: million years ago
NCBI: National Center for Biotechnology Information
TE: transposable element
WGS: whole genome sequencing
WGD: whole-genome duplication.

## Authors’ contributions

HW conducted the genome assembly, genome analysis and wrote the manuscript. VN prepared DNA and RNA for sequencing and co-wrote the manuscript. CW assisted with the genome assembly. IRS conceived and managed the project, interpreted the data and drafted the figures. ZL interpreted the data. All authors read, edited, revised, and approved the manuscript final version.

## Acknowledgements

We are grateful to the high-performance computing infrastructure provided by the University of Adelaide. We thank the Australian National Vetch Breeding programme for gifting the *V. sativa* seed.

## Additional file 2

**Figure S1:**
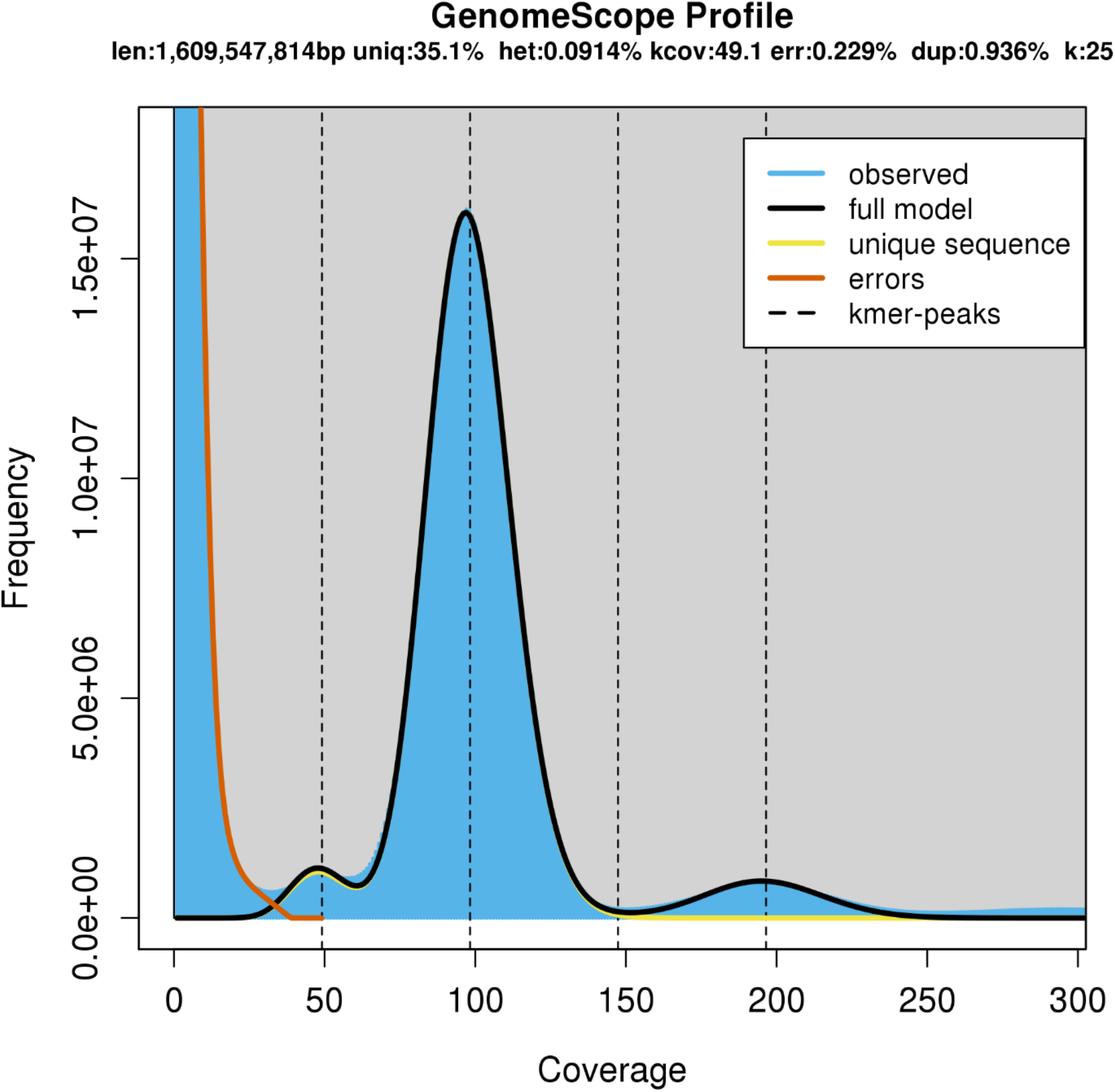
25-mer distribution of Illumina paired-end read by using GenomeScope. 25-mer occurrences (x axis) were plotted against their frequencies (y axis). Estimated genome size: 1,609,547,814 bp, estimated unique region: 35.1%, estimated heterozygosity: 0.0914%, estimated genome

**Figure S2:**
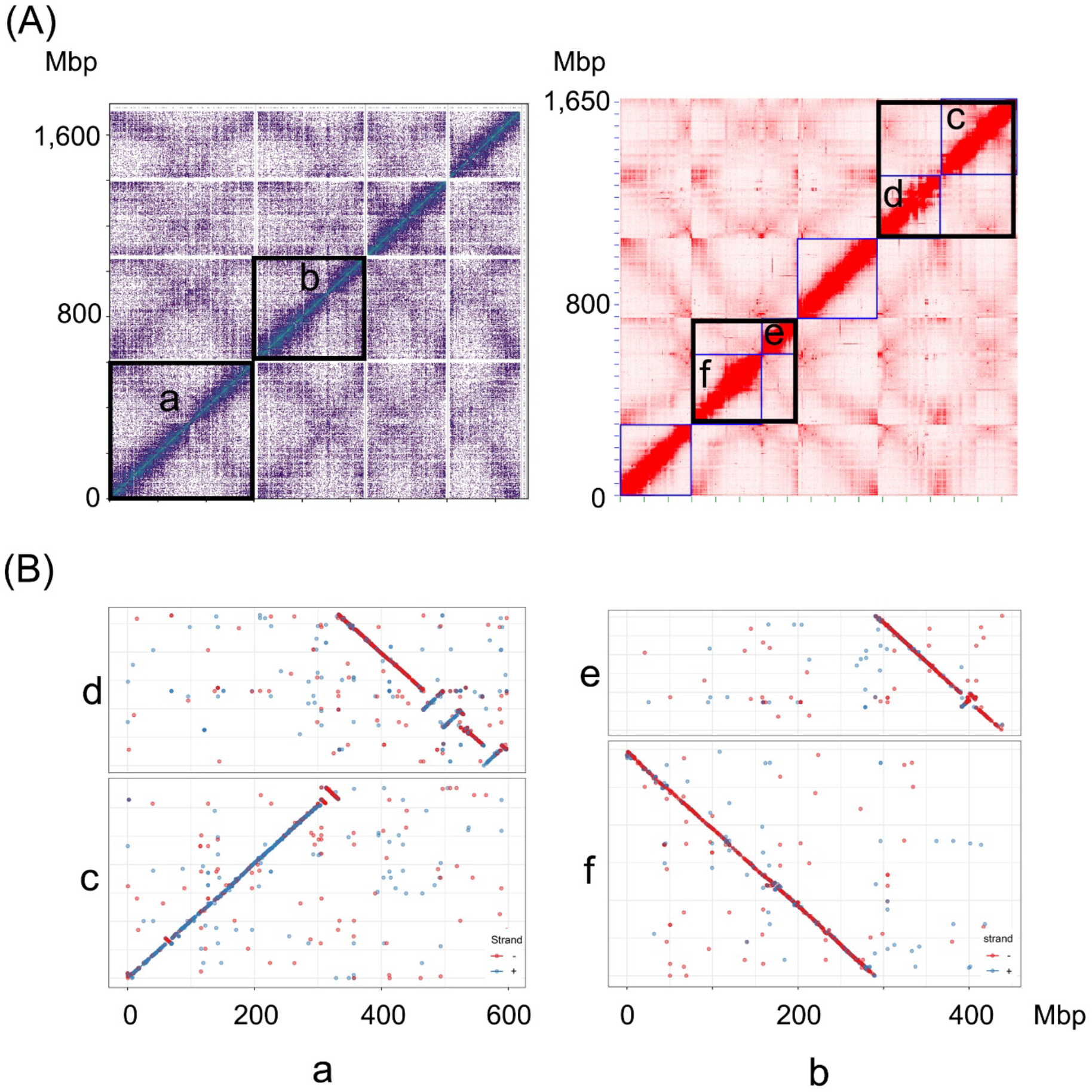
Re-solving mis-join sites in *V. sativa* pseudo-chromosomes. (A) The left figure shows the interaction heatmap of four pseudo-chromosomes generated by HiRise pipeline. Regions in black boxes show the potential mis-join pseudo-chromosomes indicated by the weak interaction signals. After re-scaffolding the genome using 3D-DNA pipeline, the mis-joins were confirmed and resulted in six pseudo-chromosomes (right figure, black boxes). (B) The collinearity between pseudo-chromosomes “a” to “c” and “d”, and between pseudo-chromosomes “b” to “e” and “f” in (A) were confirmed by the Mummer alignment.

**Figure S3:**
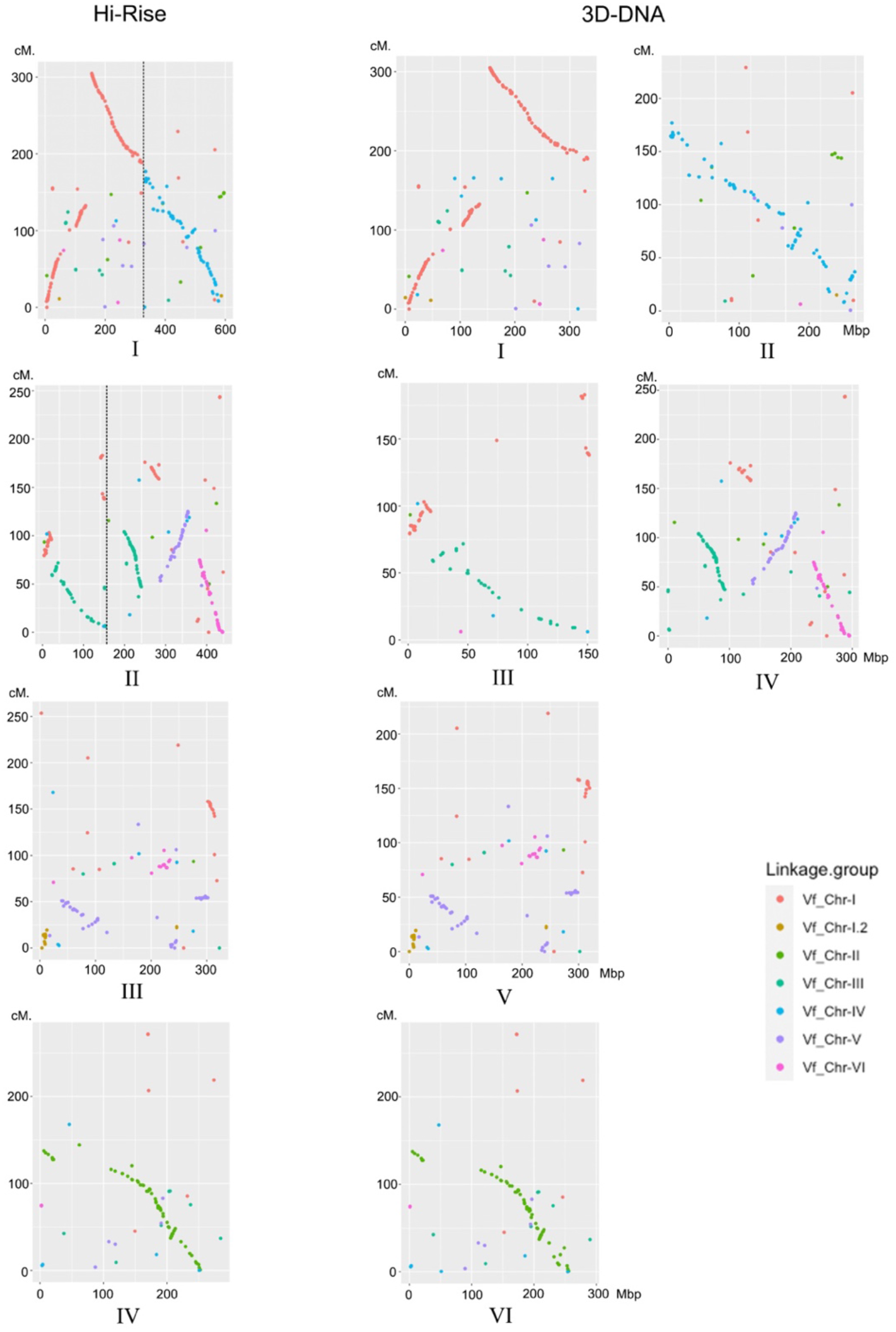
Comaprison of HiRise and 3D-DNA assembled pseudo-chromosomes to *V. faba* genetic linkage map. Left, comparison of the four HiRise pseudo-chromosomes to the genetic linkage map and right, comparison of the six 3D-DNA pseudo-chromosomes to the linkage map. The x axes present the coordinate of the pseudo-chromosomes, the y axis present the cumulative distance on the *V. faba* linkage map. Each color corresponds to a linkage group in *V. faba* linkage map. 3D-DNA pseudo-chromosomes I and II are labelled as “a” and “b” in Figure S2, respectively. HiRise pseudo-chromosomes I, II, III and IV are labelled as “c”, “d”, “e” and “f” in Figure S2, respectively.

**Figure S4:**
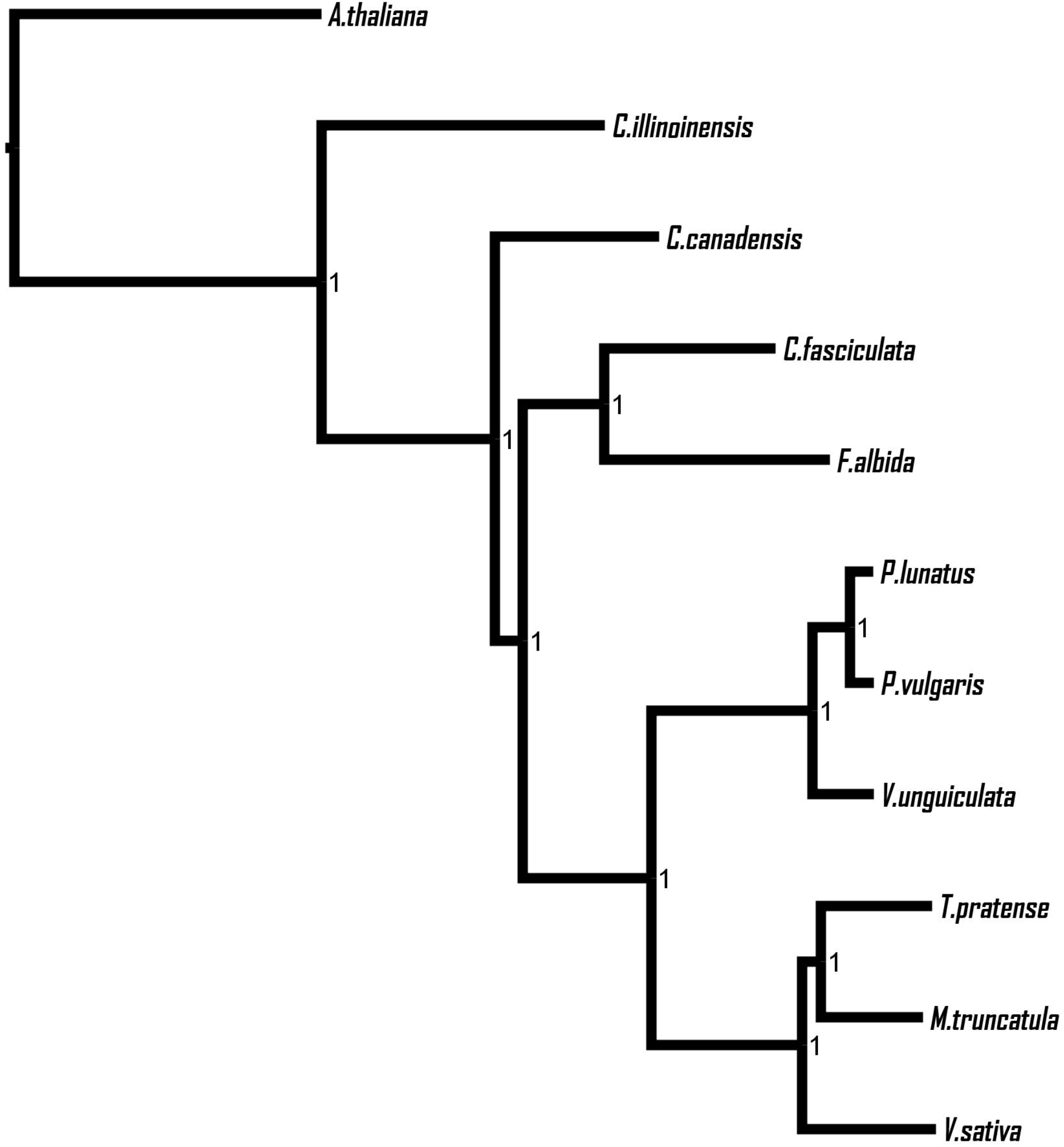
Species tree generated by Orthofinder using multiple sequence alignment approach. Node label shows the Shimodaira-Hasegawa test supporting value [1].

**Figure S5:**
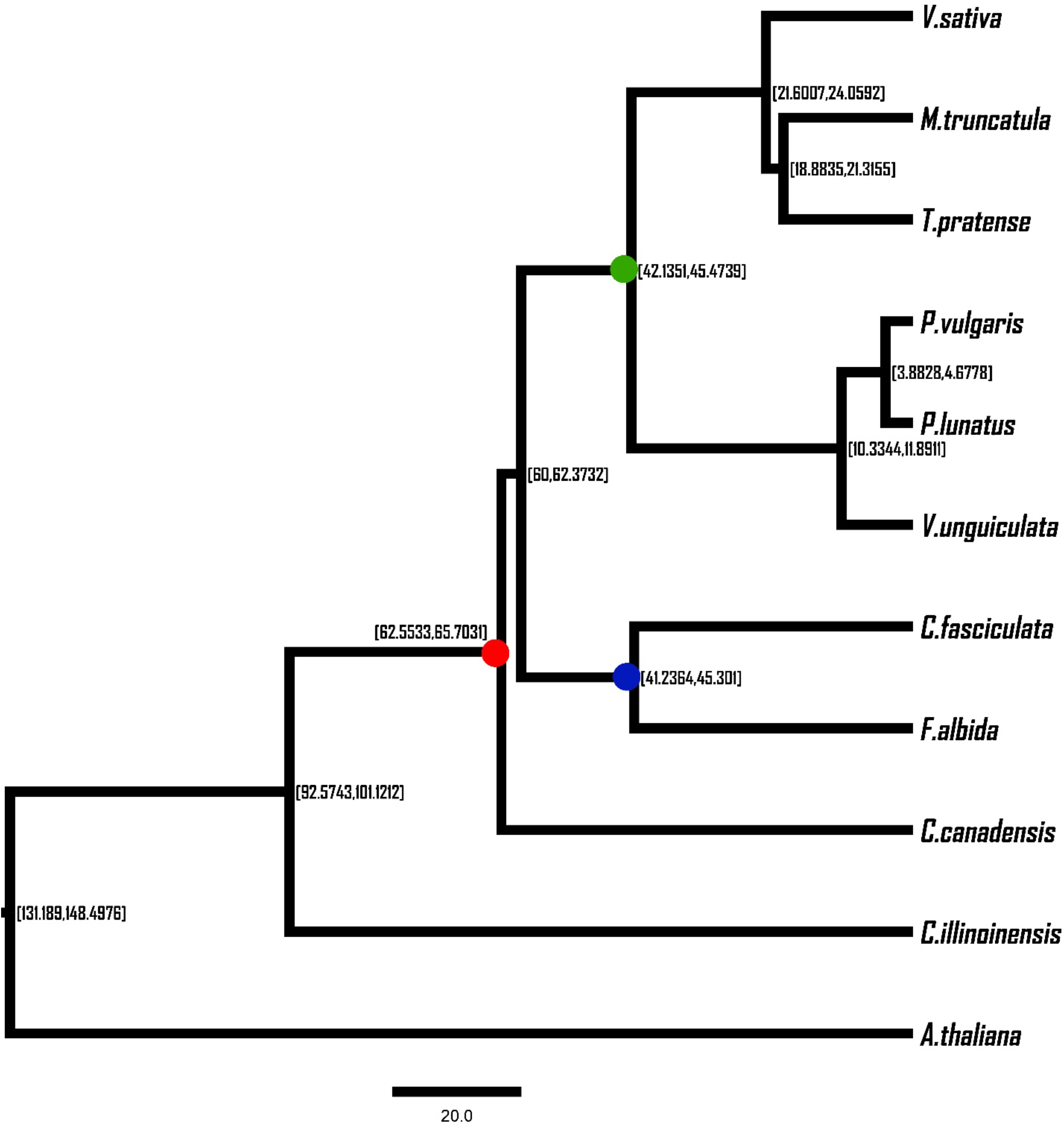
Divergence time estimation for *V. sativa* and other species. The node labels show the 95 % Highest Posterior Density of species divergence time. Unit of scale bar is million years ago. Red, blue and green dots correspond to fossil calibration points.

## References

1. Navrátilová A, Neumann P and Macas J. Karyotype analysis of four Vicia species using in situ hybridization with repetitive sequences. Annals of botany. 2003;91 7:921–6.

2. Hanelt P and Mettin D. Biosystematics of the genus Vicia L.(Leguminosae). Annual review of ecology and systematics. 1989;20 1:199–223.

3. Çceliktaş N, Can E, Hatipoğlu R and Avci S. Comparison between a wild population and cultivar of common vetch (Vicia sativa L., Fabaceae) on cytological and agronomic characteristics. 2006.

4. Zeven AC and Zhukovsky PM. Dictionary of cultivated plants and their centres of diversity: excluding ornamentals, forest trees and lower plants. Pudoc; 1975.

5. Mao Z, Fu H, Nan Z and Wan C. Fatty acid, amino acid, and mineral composition of four common vetch seeds on Qinghai-Tibetan plateau. Food chemistry. 2015;171:13–8.

6. Pfeffer M and Ressler C. β-Cyanoalanine, an inhibitor of rat liver cystathionase. Biochemical pharmacology. 1967;16 12:2299–308.

7. Meydani JA, Rahmati M, Karimi E and Aliloo AA. Dryland soil water storage susceptibility to different soil tillage practices under Vetch-Wheat crop rotation. Azarian journal of agriculture. 2014.

8. Mueller T and Thorup-Kristensen K. N-fixation of selected green manure plants in an organic crop rotation. Biological agriculture & horticulture. 2001;18 4:345–63.

9. Tenopala J, GONZÁLEZ FJ and de la Barrera E. Physiological responses of the green manure, Vicia sativa, to drought. Botanical Sciences. 2012;90 3:263–85.

10. Nguyen V, Riley S, Nagel S, Fisk I and Searle IR. Common Vetch: a drought tolerant, high protein neglected leguminous crop with potential as a sustainable food source. Frontiers in Plant Science. 2020;11:818.

11. Jia C, Dong D, Zhou Q, Searle IR and Liu Z. Significant cell differences in pod ventral suture in shatter-resistant and shatter-susceptible common vetch accessions. Crop Science. 2021;61 3:1749–59.

12. Dong R, Dong D, Luo D, Zhou Q, Chai X, Zhang J, et al. Transcriptome analyses reveal candidate pod shattering-associated genes involved in the pod ventral sutures of common vetch (Vicia sativa L.). Frontiers in plant science. 2017;8:649.

13. De la Rosa L, Zambrana E and Ramirez-Parra E. Molecular bases for drought tolerance in common vetch: designing new molecular breeding tools. BMC plant biology. 2020;20 1:1–18.

14. Sambrook J and Russell DW. Purification of nucleic acids by extraction with phenol: chloroform. Cold Spring Harbor Protocols. 2006;2006 1:pdb. prot4455.

15. Biosciences P. Guidelines for using a salt: Chloroform wash to clean up gDNA. 2019.

16. Putnam NH, O’Connell BL, Stites JC, Rice BJ, Blanchette M, Calef R, et al. Chromosome-scale shotgun assembly using an in vitro method for long-range linkage. Genome research. 2016;26 3:342–50.

17. Lieberman-Aiden E, Van Berkum NL, Williams L, Imakaev M, Ragoczy T, Telling A, et al. Comprehensive mapping of long-range interactions reveals folding principles of the human genome. science. 2009;326 5950:289–93.

18. David R, Burgess A, Parker B, Li J, Pulsford K, Sibbritt T, et al. Transcriptome-wide mapping of RNA 5-methylcytosine in Arabidopsis mRNAs and noncoding RNAs. The Plant Cell. 2017;29 3:445–60.

19. Marçais G and Kingsford C. A fast, lock-free approach for efficient parallel counting of occurrences of k-mers. Bioinformatics. 2011;27 6:764–70.

20. Vurture GW, Sedlazeck FJ, Nattestad M, Underwood CJ, Fang H, Gurtowski J, et al. GenomeScope: fast reference-free genome profiling from short reads. Bioinformatics. 2017;33 14:2202–4.

21. Macas J, Novak P, Pellicer J, Čížková J, Koblížková A, Neumann P, et al. In depth characterization of repetitive DNA in 23 plant genomes reveals sources of genome size variation in the legume tribe Fabeae. PLoS One. 2015;10 11:e0143424.

22. Koren S, Walenz BP, Berlin K, Miller JR, Bergman NH and Phillippy AM. Canu: scalable and accurate long-read assembly via adaptive k-mer weighting and repeat separation. Genome research. 2017;27 5:722–36.

23. Walker BJ, Abeel T, Shea T, Priest M, Abouelliel A, Sakthikumar S, et al. Pilon: an integrated tool for comprehensive microbial variant detection and genome assembly improvement. PloS one. 2014;9 11:e112963.

24. Simão FA, Waterhouse RM, Ioannidis P, Kriventseva EV and Zdobnov EM. BUSCO: assessing genome assembly and annotation completeness with single-copy orthologs. Bioinformatics. 2015;31 19:3210–2.

25. Van Berkum NL, Lieberman-Aiden E, Williams L, Imakaev M, Gnirke A, Mirny LA, et al. Hi-C: a method to study the three-dimensional architecture of genomes. JoVE (Journal of Visualized Experiments). 2010; 39:e1869.

26. Cairns J, Freire-Pritchett P, Wingett SW, Várnai C, Dimond A, Plagnol V, et al. CHiCAGO: robust detection of DNA looping interactions in Capture Hi-C data. Genome biology. 2016;17 1:1–17.

27. Dudchenko O, Batra SS, Omer AD, Nyquist SK, Hoeger M, Durand NC, et al. De novo assembly of the Aedes aegypti genome using Hi-C yields chromosome-length scaffolds. Science. 2017;356 6333:92–5.

28. Durand NC, Shamim MS, Machol I, Rao SS, Huntley MH, Lander ES, et al. Juicer provides a one-click system for analyzing loop-resolution Hi-C experiments. Cell systems. 2016;3 1:95–8.

29. Marçais G, Delcher AL, Phillippy AM, Coston R, Salzberg SL and Zimin A. MUMmer4: A fast and versatile genome alignment system. PLoS computational biology. 2018;14 1:e1005944.

30. Sudheesh S, Kimber R, Braich S, Forster J, Paull J and Kaur S. Construction of an integrated genetic linkage map and detection of quantitative trait loci for ascochyta blight resistance in faba bean (Vicia faba L.). Euphytica. 2019;215 3:42.

31. Ou S, Su W, Liao Y, Chougule K, Agda JR, Hellinga AJ, et al. Benchmarking transposable element annotation methods for creation of a streamlined, comprehensive pipeline. Genome biology. 2019;20 1:1–18.

32. Xu Z and Wang H. LTR_FINDER: an efficient tool for the prediction of full-length LTR retrotransposons. Nucleic acids research. 2007;35 suppl_2:W265–W8.

33. Dobin A, Davis CA, Schlesinger F, Drenkow J, Zaleski C, Jha S, et al. STAR: ultrafast universal RNA-seq aligner. Bioinformatics. 2013;29 1:15–21.

34. Brůna T, Hoff KJ, Lomsadze A, Stanke M and Borodovsky M. BRAKER2: Automatic eukaryotic genome annotation with GeneMark-EP+ and AUGUSTUS supported by a protein database. NAR genomics and bioinformatics. 2021;3 1:lqaa108.

35. Buchfink B, Xie C and Huson DH. Fast and sensitive protein alignment using DIAMOND. Nature methods. 2015;12 1:59–60.

36. Jones P, Binns D, Chang H-Y, Fraser M, Li W, McAnulla C, et al. InterProScan 5: genome-scale protein function classification. Bioinformatics. 2014;30 9:1236–40.

37. Huerta-Cepas J, Szklarczyk D, Heller D, Hernández-Plaza A, Forslund SK, Cook H, et al. eggNOG 5.0: a hierarchical, functionally and phylogenetically annotated orthology resource based on 5090 organisms and 2502 viruses. Nucleic acids research. 2019;47 D1:D309–D14.

38. Cantalapiedra CP, Hernández-Plaza A, Letunic I, Bork P and Huerta-Cepas J. eggNOG-mapper v2: Functional Annotation, Orthology Assignments, and Domain Prediction at the Metagenomic Scale. bioRxiv. 2021.

39. Schattner P, Brooks AN and Lowe TM. The tRNAscan-SE, snoscan and snoGPS web servers for the detection of tRNAs and snoRNAs. Nucleic acids research. 2005;33 suppl_2:W686–W9.

40. Nawrocki EP and Eddy SR. Infernal 1.1: 100-fold faster RNA homology searches. Bioinformatics. 2013;29 22:2933–5.

41. Kalvari I, Argasinska J, Quinones-Olvera N, Nawrocki EP, Rivas E, Eddy SR, et al. Rfam 13.0: shifting to a genome-centric resource for non-coding RNA families. Nucleic acids research. 2018;46 D1:D335–D42.

42. Emms DM and Kelly S. OrthoFinder: phylogenetic orthology inference for comparative genomics. Genome biology. 2019;20 1:1–14.

43. Ranwez V, Harispe S, Delsuc F and Douzery EJ. MACSE: Multiple Alignment of Coding SEquences accounting for frameshifts and stop codons. PloS one. 2011;6 9:e22594.

44. Capella-Gutiérrez S, Silla-Martínez JM and Gabaldón T. trimAl: a tool for automated alignment trimming in large-scale phylogenetic analyses. Bioinformatics. 2009;25 15:1972–3.

45. Bouckaert R, Vaughan TG, Barido-Sottani J, Duchêne S, Fourment M, Gavryushkina A, et al. BEAST 2.5: An advanced software platform for Bayesian evolutionary analysis. PLoS computational biology. 2019;15 4:e1006650.

46. Bouckaert RR and Drummond AJ. bModelTest: Bayesian phylogenetic site model averaging and model comparison. BMC evolutionary biology. 2017;17 1:1–11.

47. Drummond AJ, Ho SYW, Phillips MJ and Rambaut A. Relaxed phylogenetics and dating with confidence. PLoS biology. 2006;4 5:e88.

48. Heled J and Drummond AJ. Calibrated tree priors for relaxed phylogenetics and divergence time estimation. Systematic biology. 2012;61 1:138–49.

49. Magallón S, Gómez-Acevedo S, Sánchez-Reyes LL and Hernández-Hernández T. A metacalibrated time-tree documents the early rise of flowering plant phylogenetic diversity. New Phytologist. 2015;207 2:437–53.

50. Matschiner M, Musilová Z, Barth JM, Starostová Z, Salzburger W, Steel M, et al. Bayesian phylogenetic estimation of clade ages supports trans-Atlantic dispersal of cichlid fishes. Systematic biology. 2017;66 1:3–22.

51. Lyson T, Miller I, Bercovici A, Weissenburger K, Fuentes A, Clyde W, et al. Exceptional continental record of biotic recovery after the Cretaceous–Paleogene mass extinction. Science. 2019;366 6468:977–83.

52. Koenen EJ, Ojeda DI, Bakker FT, Wieringa JJ, Kidner C, Hardy OJ, et al. The origin of the legumes is a complex paleopolyploid phylogenomic tangle closely associated with the cretaceous–paleogene (K–Pg) mass extinction event. Systematic biology. 2021;70 3:508–26.

53. Wing SL, Herrera F, Jaramillo CA, Gómez-Navarro C, Wilf P and Labandeira CC. Late Paleocene fossils from the Cerrejón Formation, Colombia, are the earliest record of Neotropical rainforest. Proceedings of the National Academy of Sciences. 2009;106 44:18627–32.

54. Lavin M, Herendeen PS and Wojciechowski MF. Evolutionary rates analysis of Leguminosae implicates a rapid diversification of lineages during the tertiary. Systematic biology. 2005;54 4:575–94.

55. Fiz-Palacios O, Schneider H, Heinrichs J and Savolainen V. Diversification of land plants: insights from a family-level phylogenetic analysis. BMC evolutionary biology. 2011;11 1:1–10.

56. Silvestro D, Bacon CD, Ding W, Zhang Q, Donoghue PC, Antonelli A, et al. Fossil data support a pre-Cretaceous origin of flowering plants. Nature Ecology & Evolution. 2021;5 4:449–57.

57. De Bie T, Cristianini N, Demuth JP and Hahn MW. CAFE: a computational tool for the study of gene family evolution. Bioinformatics. 2006;22 10:1269–71.

58. Sun P, Jiao B, Yang Y, Shan L, Li T, Li X, et al. WGDI: A user-friendly toolkit for evolutionary analyses of whole-genome duplications and ancestral karyotypes. bioRxiv. 2021.

59. Pfeil B, Schlueter J, Shoemaker R and Doyle J. Placing paleopolyploidy in relation to taxon divergence: a phylogenetic analysis in legumes using 39 gene families. Systematic biology. 2005;54 3:441–54.

60. Tang H, Bowers JE, Wang X, Ming R, Alam M and Paterson AH. Synteny and collinearity in plant genomes. Science. 2008;320 5875:486–8.

## References

1. Shimodaira H. An approximately unbiased test of phylogenetic tree selection. Systematic biology. 2002;51 3:492–508.

